# Circadian desynchronization attenuates information throughput of prefrontal cortex pyramidal neurons in mice

**DOI:** 10.1101/2022.01.27.478010

**Authors:** Brandon L. Roberts, Ilia N. Karatsoreos

**Affiliations:** Neuroscience and Behavior Program, and Department of Psychological and Brain Sciences, University of Massachusetts Amherst, Amherst, MA 01003, USA

**Keywords:** prefrontal cortex, circadian rhythms, pyramidal neurons, GIRK, circadian disruption

## Abstract

The prefrontal cortex (PFC) is heavily involved in cognitive and emotional processes, including working memory, cognition, stress responses, and fear associated behaviors. Many PFC-associated behaviors are time-of-day dependent, and disruption of daily rhythms negatively impacts these behavioral outputs, yet how the disruption of daily rhythms impacts the fundamental function of PFC neurons, and the mechanism(s) by which this occurs, remains unknown. Using a mouse model, we demonstrate that the activity and action potential dynamics of prelimbic PFC neurons are regulated by time-of-day in a sex specific manner. Further, we show that postsynaptic K^+^ channels play a central role in mediating these rhythms, suggesting an intrinsic gating mechanism mediating information throughput. Finally, we demonstrate that environmental circadian desynchronization alters the intrinsic functioning of these neurons in part by increasing sensitivity GIRK channel activation. These key discoveries demonstrate daily rhythms contribute to the mechanisms underlying the essential physiology of PFC circuits, and provide potential mechanisms by which circadian disruption may impact the fundamental properties of neurons.

**Significance Statement:** Disruption of circadian rhythms, such as shift work and jet lag, are associated with negative physiological and behavioral outcomes, including changes in affective state, cognitive function, learning and memory. The prefrontal cortex (PFC) plays a critical role in these functions, yet how daily rhythms and desynchronization of these rhythms impact the physiology of neurons in the PFC is unknown. Here we demonstrate that daily rhythms impact the physiological function of PFC neurons in a sex-dependent manner, and that environmental circadian desynchronization alters PFC function irrespective of time-of-day. These findings provide not only a physiological context to the neural and behavioral changes associated with circadian desynchronization, but also highlight the importance of considering the temporal dimension in studies of neural circuits.

## Introduction

Circadian (daily) rhythms are ubiquitous in nature and play a central role in health and disease. Synchrony between internal clocks and external time are important for optimal organismal function, while their desynchronization leads to myriad negative mental and physical health outcomes. Understanding the functional relevance of normal circadian rhythms, or their desynchronization, in neural function is important if we are to fully appreciate how disrupted clocks impact health outcomes.

The prefrontal cortex (PFC) serves as a critical node in cognition, emotional systems involved in fear learning and extinction, stress responses, and learning and memory, all of which vary over the course of the day (1–5). The PFC is comprised of an array of cell types, including excitatory pyramidal neurons and inhibitory interneurons, which together impact behavior by relaying information to other brain regions including the amygdala and hippocampus, both of which also demonstrate circadian rhythms (6–9). In mouse, the prelimbic area (pl) of the PFC is divided into six distinct layers, each with distinct inputs and projections. Specifically, neurons in layer 2/3 play a major role in working memory and behavioral plasticity and is involved in stress and depressive behaviors (7, 10–12). Our previous work shows that environmental circadian desynchronization has profound impacts on behavioral flexibility, responses to novel environments, and on the morphology of these layer 2/3 plPFC pyramidal neurons (13). Yet the impacts of circadian desynchronization on neuronal function remains unknown.

The majority of PFC pyramidal neurons are intrinsically quiescent in their resting state and regulate information throughput via a wide array of ion channels, such as cyclic-nucleotide-gated non-selective cation (HCN) channels, sodium (Na^+^), calcium (Ca^2+^) and potassium (K^+^) channels, including g-protein inward rectifying K^+^ (GIRK) channels that mediate postsynaptic throughput of synaptic currents (11, 14–16). In the suprachiasmatic nucleus (SCN) brain clock, changes in sodium (Na^+^), K^+^, and Ca^2+^ ion channel function mediate daily rhythms in the spontaneous activity, and action potential dynamics of neurons (17, 18). How these channels might impact daily rhythms in PFC function and the gating of information throughput remains an important gap in our knowledge.

In the present set of studies, we show that circadian desynchronization markedly alters the intrinsic function of plPFC pyramidal neurons. Second, we demonstrate that time-of-day clearly drives changes in the fundamental neurophysiological properties of these neurons, including information throughput, defined here as a combination of resting membrane potential, action potential threshold, and action potential firing rate after current injection. Third, we identify that a potential mechanism for these daily changes in the plPFC involves rhythms in K^+^ channel function. Finally, we demonstrate that one mechanism by which environmental circadian desynchronization decreases information throughput in these neurons is by altering GIRK channel function. Our findings reveal that time-of-day is a critical factor that significantly changes neurophysiological properties of plPFC pyramidal neurons and that environmental circadian desynchronization has sizeable consequences for neural function independent of time of day. Together, this work further establishes these cells as an important substrate for some of the negative outcomes that accompany circadian desynchronization.

## Results

### Circadian disruption decreases excitability of PFC pyramidal neurons

The regional and cell-specific heterogeneity in electrophysiological properties of PFC pyramidal neurons has been described in multiple species(11, 19, 20). Here we define type I pyramidal neurons as those that are quiescent (non-firing) at rest, slow-firing after stimulation, have a longer half-width, and shallow antipeak amplitude, while all other phenotypes are considered type II. We focused the present studies on the type I subset, which composed ∼80% of recorded neurons (***SI Appendix*, Fig. S2A-H**). The type I subset of layer 2/3 pyramidal neurons of the plPFC were identified visually by anatomical location and phenotypically by electrophysiological characterization in *ex vivo* coronal brain slices (**Fig. 1A, B;** *left*, ***SI Appendix*, Fig. S1A-D, Fig. S2A-H**). Pyramidal neurons were identified by shape and lucifer yellow (LY; 0.2%) was added to the patch pipette for confirmation of an apical dendrite (**Fig. 1B;** *right*).

**Figure 1.**
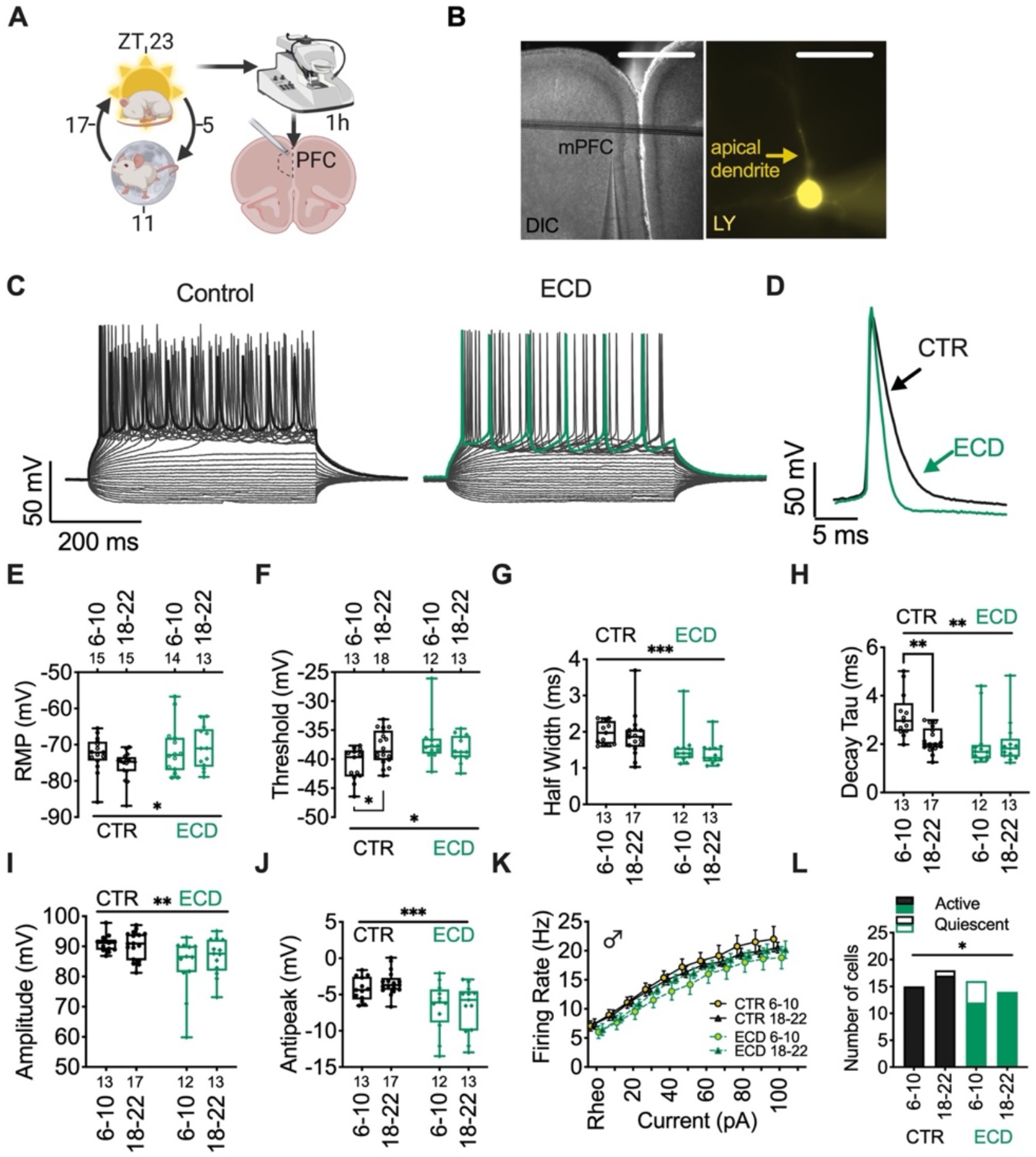
Environmental Circadian Desynchronization decreases information throughput and alters fundamental properties of PFC pyramidal neurons independent of time-of-day. **(A)** Diagram of methodological approach for ZT times (*left*) of *ex vivo* slice collection (*right*) **(B)** mPFC slice (*left;* scale 1mm) and layer 2/3 pyramidal neuron backfilled with lucifer yellow (LY, *right;* scale 10 μm). **(C)** Representative trace of current step recording with maximal AP firing and **(D)** individual evoked APs highlighted at ZT6-10 in control (CTR) (*left; black*) and ECD (*right; bluish green*) in male mice. **(E)** membrane potential (Time: F (1, 52) = 0.9617; *p* = 0.33, Treatment: F (1, 52) = 4.275, *p* = 0.04), **(F)** action potential firing threshold (Time: F (1, 52) = 1.1143; *p* = 0.29, Treatment: F (1, 52) = 4.867; *p* = 0.03), **(G)** half width (Time: F (1, 51) = 0.8106; *p* = 0.37, Treatment: F (1, 51) = 13.80; *p* < 0.001), **(H)** decay tau (Time: F (1, 51) = 4.705; *p* = 0.0348, Treatment: F (1, 51) = 7.294; *p* = 0.0094), **(I)** amplitude (Time: F (1, 51) = 0.4232; *p* = 0.52, Treatment: F (1, 51) = 10.24; *p* = 0.002), and **(J)** action potential antipeak (Time: F (1, 51) = 0.292; *p* = 0.5913, Treatment: F (1, 51) = 17.70; *p* = 0.0001) at ZT 6-10 and 18-22 in male mice. **(K)** Firing rate (F (3, 54) = 0.832; *p* = 0.48), Treatment: F (1, 53) = 1.081; *p* = 0.30) and **(L)** number of active vs quiescent neurons after 200pA current injection. Box plots represent median, min/max, and second and third quartiles. N-values for number of cells inset on x-axis. Two-way ANOVA for main effects and interaction, with Šídák’s multiple comparisons post-hoc analysis for ZT bin. *\*p <0*.*05, **p <0*.*01, ***p <0*.*001*.

Environmental circadian desynchronization (ECD), implemented by housing in a 10:10h light:dark cycle, negatively impacts cognitive flexibility and alters both morphology and dendritic connections of layer 2/3 plPFC pyramidal neurons in mice (13). We hypothesized that the impact of ECD on these neurons would extend to their endogenous physiological properties. To test this, we used whole-cell patch clamp techniques and measured changes in resting state properties and action potential dynamics at zeitgeber times (ZT) 6-10 and 18-22 (calculated as 50min/h in ECD mice; **Fig. 1E-J**). We observed a main effect of ECD, independent of time-of-day, on the resting membrane potential and action potential firing threshold (**Fig. 1E, F**). Further, ECD had a main effect on all components of action potential dynamics that were measured, including half width, decay tau, amplitude, and antipeak (**Fig. 1C,D,G-J**). Finally, all other effects of time observed in control mice, such as action potential threshold and decay tau, was lost in ECD mice. While there was no ECD effect on evoked action potential firing rate (as calculated from rheobase) after current injection (**Fig. 1K-L**), we did not include neurons that failed to elicit action potentials after current injections. In a separate Chi-squared analysis of active vs quiescent neurons, we observed a significant increase in the number of quiescent neurons in ECD mice (**Fig. 1L**). Together these data show that independent of time-of-day, ECD alters the fundamental intrinsic properties of plPFC pyramidal neurons.

### Resting membrane potential of prelimbic layer 2/3 pyramidal neurons is rhythmic in male mice

In pursuit of the mechanistic underpinnings for how ECD changes information throughput in the PFC, we first needed to define how daily rhythms impact the normal function of these neurons. To test our hypothesis that time-of-day impacts the basal electrophysiological properties of pyramidal neurons, we used whole-cell patch clamp techniques and measured resting membrane potential (RMP) and membrane resistance (Rm) at ZT bins: 0-4, 6-10, 12-16, and 18-22 in male (*n* = 6 mice/group) and female mice (*n* = 4-6 mice/group) (**Fig. 2A-C**). There was a main effect of time on RMP (no effect of sex) and notably, within group post-hoc analysis revealed that plPFC pyramidal neurons are more depolarized at ZT6-10 (light period) in male mice, when compared to 12-16 and 18-22 (dark period). Post-hoc analysis did not reveal a time-of-day effect on RMP in female mice (**Fig. 1B**). We did not observe an interaction between sex and time in any of our measures; however, there was a main effect of sex on Rm (**Fig. 1C**). Together, these data demonstrate that the RMP of plPFC pyramidal neurons in male mice changes throughout the light/dark (LD) cycle.

**Figure 2.**
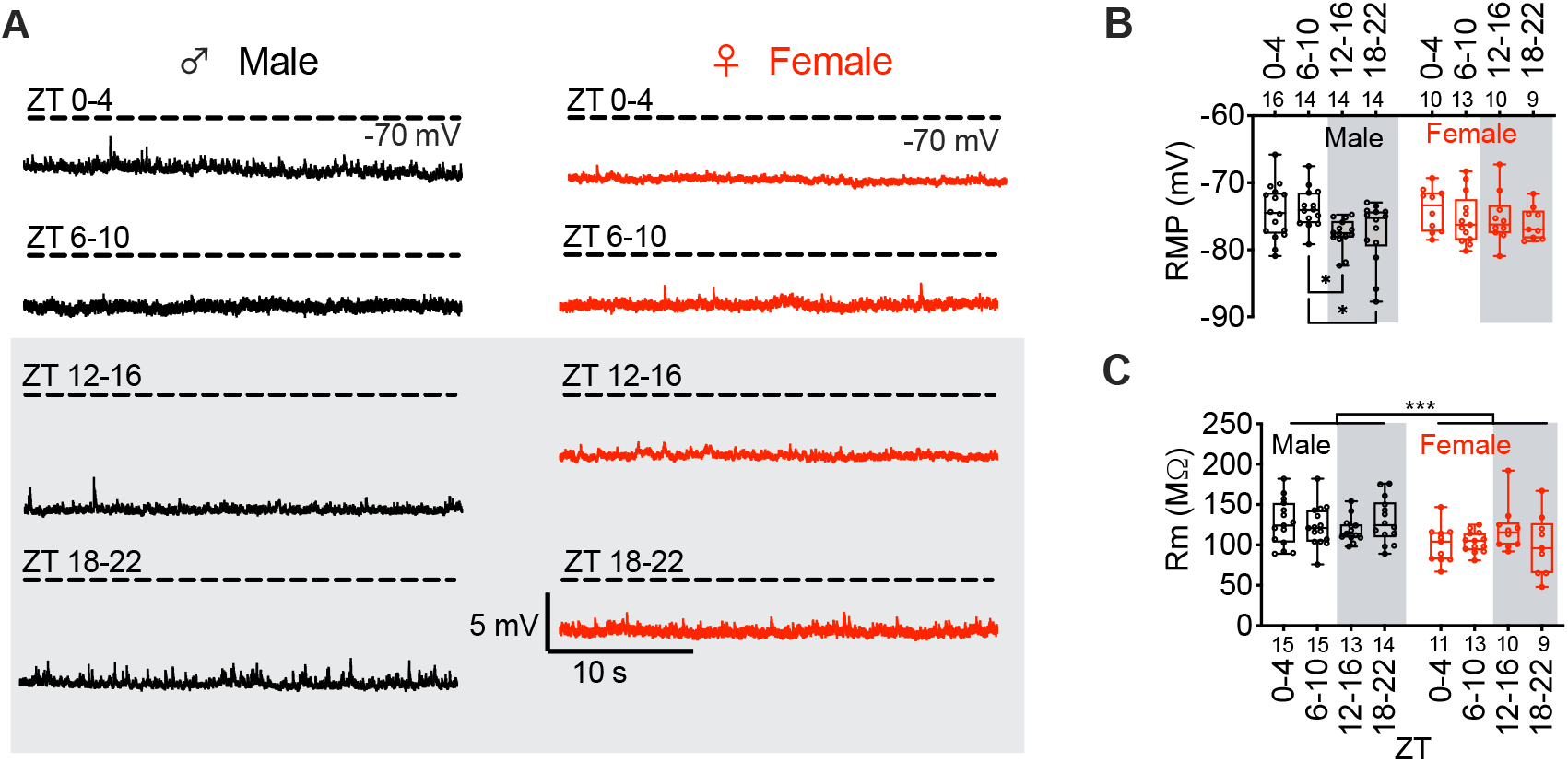
Resting membrane potential of layer 2/3 mPFC pyramidal neurons is rhythmic in male mice. **(A)** Representative traces of current clamp recordings from male and **(D)** female mice at each ZT bin. **(B)** Mean and individual data points for membrane potential (RMP) at ZT0-4, 6-10, 12-16, and 18-22 in male (*black; left*) and female (*red; right*) mice (Time: F (3, 92) = 3.126; *p* = 0.03, sex: F (1, 92) = 0.718; *p* = 0.40) **(C)** Mean membrane resistance (Rm; Time: F (3, 92) = 0.250; *p* = 0.86, Sex: F (1, 92) = 13.25; *p* = <0.001) binned by ZT. Box plots represent median, min/max, and second and third quartiles. N-values for number of cells inset on x-axis. Two-way ANOVA for main effects and interaction with a within group Tukey post-hoc analysis for ZT bin, * *p* < 0.05, \*\*\**p* < 0.001.

### Action potential throughput is decreased during the active period in plPFC pyramidal neurons

To understand the functional implications of daily rhythms in RMP and postsynaptic ion channel function for information throughput, we tested how time-of-day impacts action potential dynamics. We utilized a 10-pA current injection protocol to evoke action potentials at ZT0-4, 6-10, 12-16, and 18-22 and observed a main effect of time for membrane potential threshold of action potential firing, with post-hoc analysis revealing an increased threshold for firing late in the dark period (ZT18-22) when compared to ZT6-10 in male mice (*n* = 6 mice/group) (**Fig. 3A-C**). Consistent with the null effect of time on RMP in female mice, there was no effect of time on action potential threshold (*n* = 4-6 mice/group) (**Fig. 3C**). Further, although there was no effect on amount of current needed to elicit an action potential (rheobase) in male mice, once rheobase was reached, subsequent current injections evoked action potential firing at a lower frequency during the dark period (**Fig. 3D, E**). Although we did not observe a time-of-day effect on action potential amplitude or half-width in male mice (**Fig. 3F-G**), decay tau, a component of action potential firing that is modulated to a large extent by K^+^ channels, was reduced between ZT6-10 and 18-22 during our initial experiments (**Fig. 1H**) and likely failed to reach significance in this experiment do to the interaction between sex and time (**Fig. 3H**), as the values for these data in control males are highly consistent between experiments. Together, these data suggest that plPFC pyramidal neurons are not only more hyperpolarized during the light period, but are functionally more difficult to activate, requiring larger depolarizations to elicit action potentials and relay information downstream.

**Figure 3.**
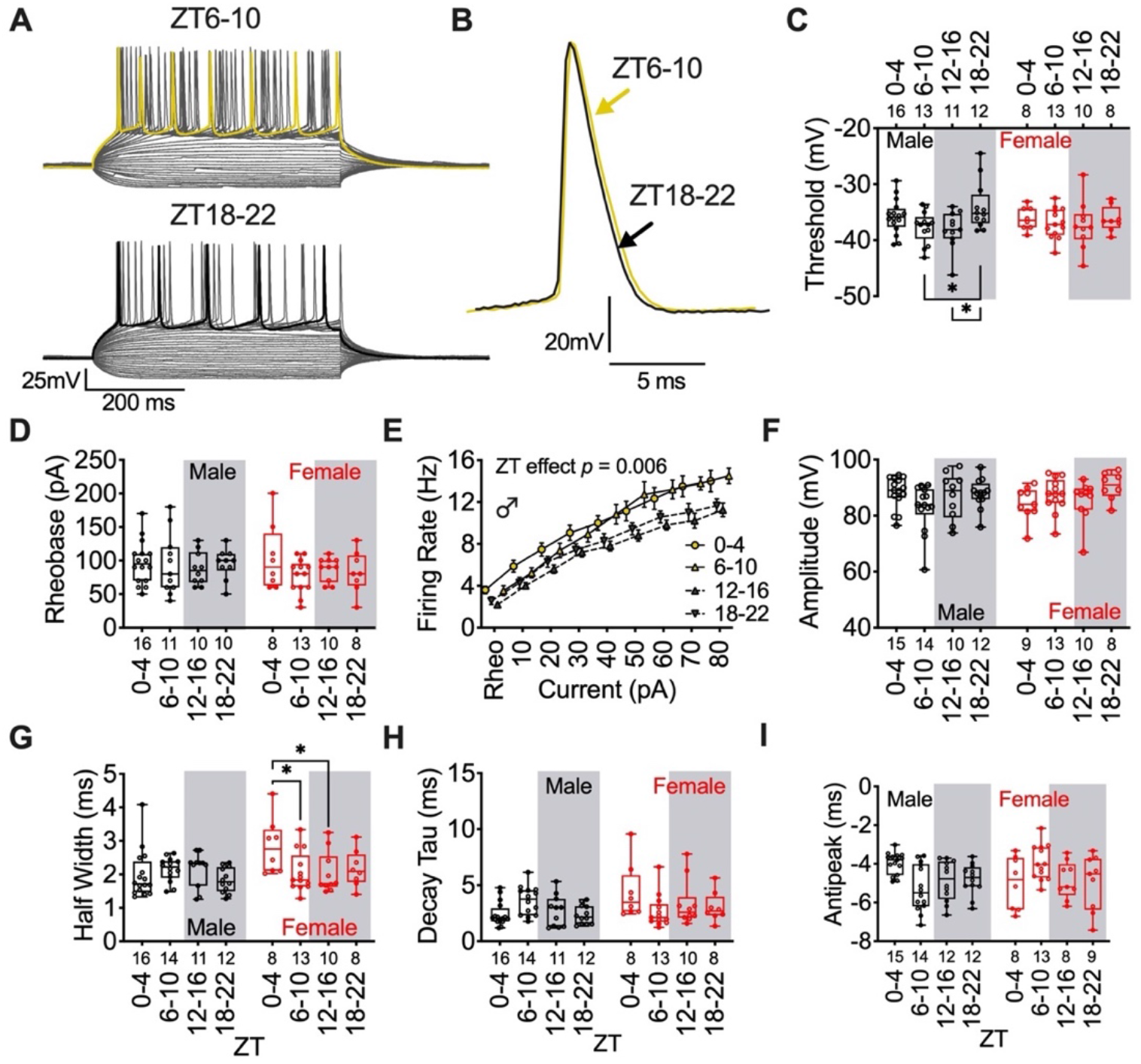
Information throughput is decreased during the active period in male mice. **(A)** Representative trace of current step recording with maximal AP firing highlighted at ZT6-10 (*top; yellow*) and 18-22 (*bottom; black*) and **(B)** individual evoked APs in male mice. **(C)** Mean AP threshold (Time: F (3, 83) = 2.809; *p* = 0.04, Sex: F (1, 83) = 0.1388; *p* = 0.71) **(D)** Rheobase (Time: F (3, 78) = 0.9519; *p* = 0.42, Sex: F (1, 78) = 0.6786; *p* = 0.41), **(E)** evoked firing rate (from rheobase; male mice Time: F (3, 41) = 4.815; *p* = 0.006), **(F)** amplitude (Time: F (3, 83) = 1.489; *p* = 0.22, Sex: F (1, 83) = 0.1132; *p* = 0.74), **(G)** half width (F (3, 84) = 1.993; *p* = 0.12, Sex: F (1, 84) = 4.045; *p* = 0.049), **(H)** decay tau (Time: F (3, 84) = 0.7564; *p* = 0.5217, Sex: F (1, 84) = 3.442; *p* = 0.0671) and **(I)** antipeak (Time: F (3, 84) = 0.8137; *p* = 0.49, Sex: F (1, 84) = 0.0505; *p* = 0.82) at each ZT bin in male and female mice. Box plots represent median, min/max, and second and third quartiles. Error bars on I-V plots represent ± SEM. N-values for number of cells inset on x-axis. Two-way ANOVA for main effects and interaction, with a within group Tukey post-hoc analysis for ZT bin and/or current injection. *\*p <0*.*05*.

### K^+^ channel activity regulates postsynaptic rhythmic activity in pyramidal neurons of male mice

Given that only male mice display daily rhythms in RMP, we hypothesized that changes in ion channel properties may play a role in setting the functional tone of plPFC pyramidal neurons specifically in male mice. Changes in ion channel properties can be assessed by changes in current density and cellular conductance as measured by the current-voltage (I-V) relationship. To investigate daily changes in the I-V relationship, we used a potassium (K^+^) gluconate internal solution and performed an inactivation protocol in which neurons held at −70mV were stepped from 30mV to −120mV, with subsequent −10mV sweeps. (**Fig. 4A, B**). At each ZT bin (*n* = 6 male mice/group), we analyzed the steady-state current density (current normalized to cell capacitance) of the inward (−120 mV to −70mV steps (*K1*); **Fig. 4A-C**) and outward current (0 mV to 30mV steps (*K2*); **Fig. 4A, B, D**), as well as daily changes in *K1* and *K2* conductance (*g*) normalized to cell capacitance (**Fig. 4I, J**). The I-V relationship demonstrated a clear increase in inward current at hyperpolarized holding voltages and an increase in delayed outward rectifying current at depolarized voltages early in the dark period (ZT12-16) (**Fig. 4A-D**). This effect translated into increased cell conductance late in the light period and early in the dark period, or near the transition to lights off (**Fig. 4I, J)**.

**Figure 4.**
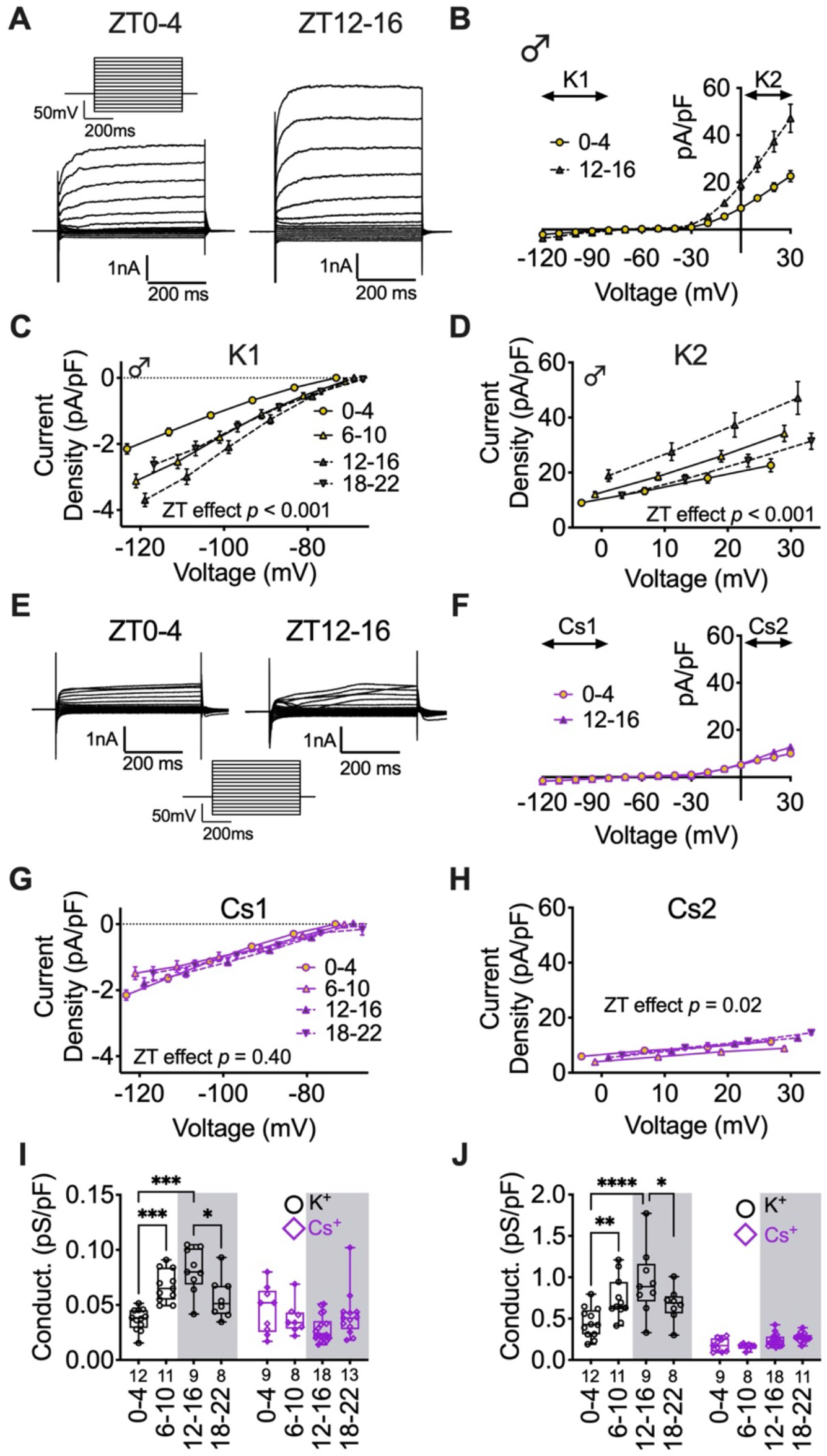
K^+^ channels are required for time-of-day effects on membrane conductance. **(A)** Voltage-step protocol (*top left*) and representative voltage-step traces of I-V relationship at ZT 0-4 (*left*) and 12-16 (*right*) in male mice. **(B)** Averaged I-V relationship for ZT0-4 and 12-16 (*normalized to cell capacitance*) with a K^+^ internal solution. **(C)** Current density of K1 (Time: F (3, 37) = 9.115; *p* = <0.001) and **(D)** K2 (Time: F (3, 37) = 9.055; *p* = <0.001) I-V relationships at ZT0-4, 6-10, 12-16, and 18-22. **(E)** Representative voltage-step traces of **(F)** I-V relationship at ZT 0-4 (*left*) and 12-16 (*right*) with a Cs+ internal solution. **(G)** Current density of Cs1 (Time: F (3, 48) = 1.009; *p* = 0.40) and **(H)** Cs2 (Time: F (3, 45) = 3.540; *p* = 0.02) I-V relationships at each ZT bin. **(I)** Comparison each ZT bin for K1 and Cs1 (Time: F (3, 80) = 2.682; *p* = 0.05, Internal: F (1, 80) = 37.22; *p* = <0.001), and **(J)** K2 and Cs2 (Time: F (3, 81) = 7.896; *p* = 0.0001, Internal: F (1, 81) = 141.1; *p* = <0.0001) normalized cell conductance. Two-way ANOVA for main effects and interaction, with a within group Tukey post-hoc analysis for ZT bin, voltage, and internal solution. Box plots represent median, min/max, and second and third quartiles. Error bars on I-V plots represent ± SEM. N-values for number of cells inset on x-axis. \**p* < 0.05, \*\**p* < 0.01, \*\*\**p* < 0.001, \*\*\*\**p* < 0.0001.

Since ionic conductance was highest at ZT12-16, the same ZT bin that RMP was most hyperpolarized (**Fig.2B**), we predicted that this increased conductance was due to increased K^+^ channel activity. To determine if daily changes in current density and cell conductance was dependent on K^+^ channel activity, we utilized a K^+^ free Cs^+^-based internal recording solution to block K^+^ currents (*n* = 6-8 male mice/group). This preparation completely abolished the time-of-day effect on inward currents and greatly reduced the time-of-day effect on outward currents (**Fig. 4E-H**). Further, the time-of-day effect on cell conductance was blocked by Cs^+^ (**Fig. 4I,J**). Of particular note, blockade of inward K^+^ currents via internal Cs^+^ appeared to have little effect at ZT0-4, suggesting minimal K^+^ channel conductance at this ZT bin (**Fig. 4C,G**). Together, these data demonstrate that K^+^ channels contribute to daily rhythms in the cellular conductance of plPFC pyramidal neurons.

### Circadian disruption upregulates GIRK channel activation independent of daily rhythms

Our previous work showed that ECD impacts the PFC by reducing pyramidal cell dendritic arborizations and impairs cognitive flexibility (13). Notably, G protein-gated inwardly rectifying K^+^ (GIRK) channels in the PFC contribute to the regulation of cognitive flexibility and contribute to action potential firing threshold (21–23). To test if increased sensitivity to GIRK channel activation serves as a potential mechanism for decreased information throughput in ECD mice we bath applied the selective GIRK channel agonist ML297 (10 μM)(23, 24) and measured changes in holding current (**Fig. 5A, B**). To specifically measure the chronic impact of ECD on pyramidal neuron function, we combined recordings from ZT6-10 and 18-22 to eliminate time-of-day as an independent variable. We predicted that application of a GIRK agonist would induce larger outward currents of PFC pyramidal neurons in ECD mice. maximal GIRK channel activity. Compared to control mice, ML297 induced outward currents were greater in ECD mice. We predicted that this increase in outward current would translate to decreased information throughput, or the decreased firing rate of pyramidal neurons, for equal injections of current. We evoked action potentials before and after the bath application of the selective GIRK channel agonist ML297 (**Fig. 5C**). Compared to control mice, most neurons recorded from ECD mice failed to elicit action potentials in the presence of ML297 irrespective of the light/dark period, suggesting the impact of GIRK channels on cellular function and inhibition of information throughput is more robust in ECD mice (**Fig. 5D**). A small subset of neurons in CTR and ECD mice did not fire in control aCSF conditions after the maximal current injection for our stimulation protocol and these neurons were removed from this portion of analysis. We plotted the firing rate of neurons that fired in control conditions and observed a robust decrease in action potential firing for CTR and ECD groups (**Fig. 5E,F**). The firing rate after ML297 application approached significance for the main effect of CTR vs ECD conditions. This lack of significance is likely due to the very low firing rate in the presence of ML297, but did reach significance when plotted as a percent reduction in firing rate after ML297 application. Together, these data demonstrate that GIRK channel activity contributes to changes in action potential throughput induced by ECD.

**Figure 5.**
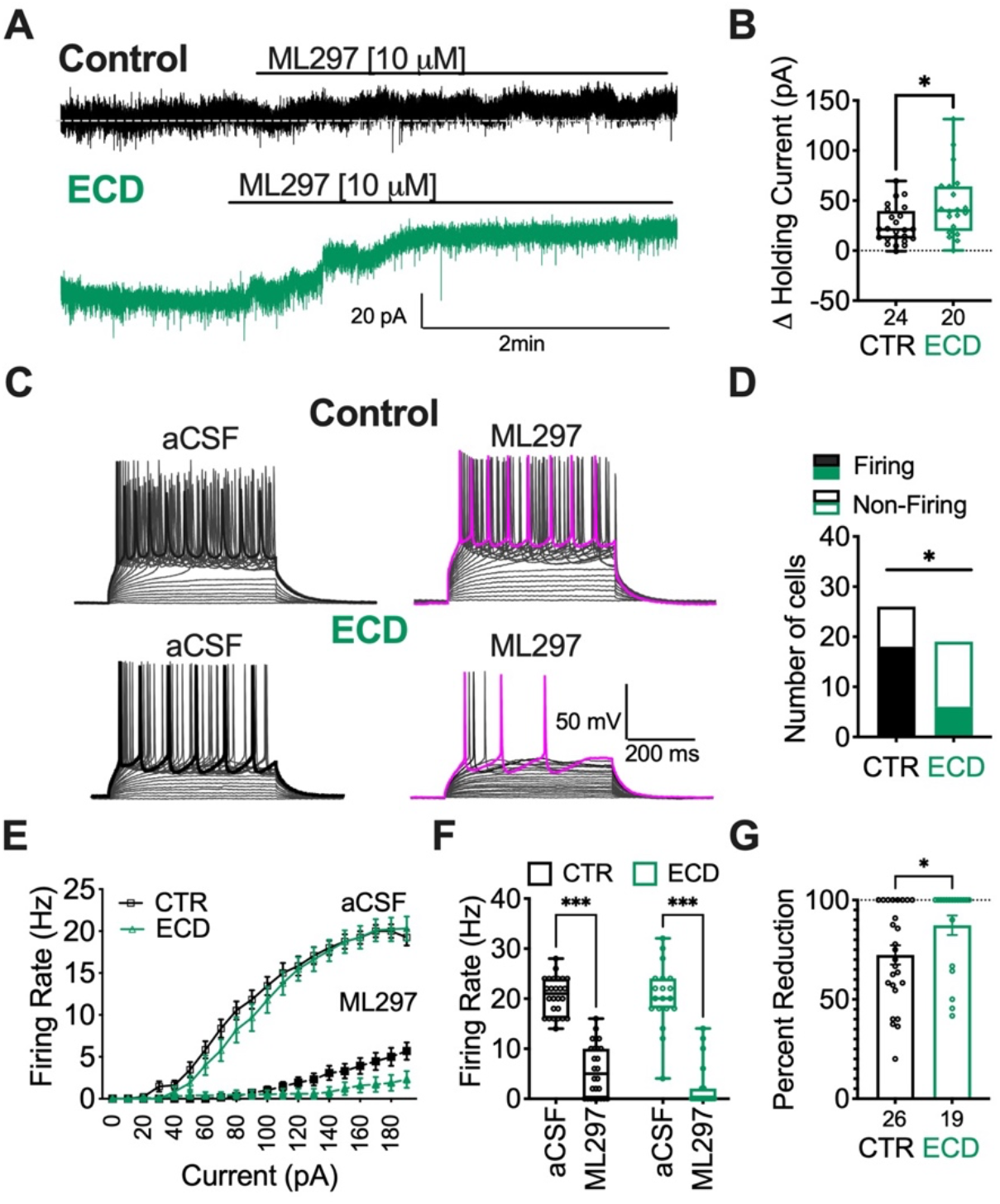
Circadian disruption increases sensitivity to GIRK channel activation in PFC pyramidal neurons. **(A)** Representative voltage clamp trace of holding current before and after application of ML297 (10 μM) and **(B)** mean change in holding current after application of ML in neurons from CTR and ECD mice (t = 2.695, df = 42, *p* = 0.01). **(C)** Representative trace of current step recording with maximal AP firing highlighted in control (CTR; *top*) and ECD (*bottom*) before (*left*) and after (*right*) application of ML297 (10 μM) in male mice. **(D)** Number of cells that elicited action potentials in the presence of ML297 (Chi-square = 6.253, df = 1; *p* = 0.012). **(E)** Average firing rate before and after treatment with ML297 at subsequent 10pA current steps, **(F)** maximal firing rate (Drug: F (1, 43) = 276.6; *p* < 0.001, Treatment: F (1, 43) = 2.10; *p* = 0.16), and **(G)** percent reduction in firing rate after application of ML297 in CTR and ECD mice (t = 2.101, df = 43, *p* = 0.042). Error bars represent ± SEM. N-values for number of cells inset on x-axis. *\*p <0*.*05, ***p <0*.*001*.

## Discussion

While there have been several well executed studies that demonstrate the presence of daily rhythms in neurophysiological function in the hippocampus and brainstem, none have included the PFC, and many have largely focused on extracellular field recordings (25–29). Thus, previous studies have not shed light on how circadian desynchronization impacts the fundamental electrophysiological processes in higher brain areas. In this study we present five main findings. First, we illustrate that circadian desynchronization disrupts multiple properties of neural function and reduces information throughput layer 2/3 plPFC pyramidal neurons independent of time-of-day. Second, we demonstrate that in male mice, these neurons are hyperpolarized during the early portion of the dark period when compared to the latter portion of the light period. Third, we demonstrate that male mice display distinct changes in ion channel activity and action potential kinetics, showing increased action potential firing threshold and decreased decay tau during portions of the dark period. Fourth, we identify that changes in K^+^ channel activity likely serves as a potential mechanism underlying time-of-day changes in the RMP and action potential firing rates of plPFC pyramidal neurons. Finally, we illustrate that circadian desynchronization induces an increased sensitivity to GIRK channel activation, which serves as a potential mechanism contributing to altered neural activity in the PFC of ECD mice. By identifying the intrinsic properties and synaptic inputs of plPFC pyramidal neurons, and a putative cellular substrate for the effects of circadian desynchronization, these findings allow us to better understand the relationship between circadian function/dysfunction, intrinsic PFC circuitry and its outputs.

Changes in PFC function underly numerous psychiatric disorders including bipolar, post-traumatic stress disorder (PTSD), attention deficit disorder, and deficits in learning and memory (3–5, 30). There is growing evidence of links between circadian rhythms and both PFC function and many of these behavioral conditions (1, 31–34). Previous work from our group has demonstrated that extracellular lactate (a functional output of neural metabolism) shows circadian rhythms in the medial (m) PFC, and that ECD alters the morphology of mPFC neurons and PFC mediated behaviors (13, 35). The present studies contextualize these results, and are the first to directly test whether cell autonomous activity and synaptic inputs onto PFC neurons are rhythmic, define contributing mechanisms by which this occurs, and identify how circadian disruption impacts these functions.

Our finding that the resting state of plPFC neurons is more hyperpolarized during the dark period, when nocturnal mice are awake and active, suggests that these neurons are more quiescent and require a higher degree of information input (i.e. synaptic activation) before eliciting a response and sending downstream signals to other brain regions. That is, there is increased gating in this circuit at night. On the surface, it seems counter-intuitive that plPFC pyramidal neurons would show more gating during the dark (active) phase, than the light (rest) phase. A functional hypothesis for this finding is that stronger gating during the active phase serves as a mechanism for selective information throughput in response to environmental stimuli. Information filtering is paramount for proper behavioral output, and too low of a threshold may result in overactivation as the animal engages with its environment. For example, pharmacological studies have demonstrated that activation of the plPFC with neurotensin agonists or the sodium channel activator veratrine lead to anxiogenic behaviors, likely through increased glutamate release (36–38). Behaviorally, autism spectrum disorder mouse models, with hyperactive ventral hippocampus to mPFC excitatory inputs, display impaired social memory that is rescued by inhibition of this circuitry, supporting the notion that overactivation of these neurons may be detrimental to appropriate input-output relationships and their impact on behavior (39).

To understand the functional relevance of cell endogenous changes in resting state, we investigated how time-of-day affects action potential dynamics, since action potential firing is a functional measure for information throughput. Notably, this measure changes with time-of-day in the hippocampus, and in response to sleep deprivation in the PFC (40, 41). Action potentials are dependent on voltage-gated ion channels, and changes in K^+^ channel activity can alter action potential firing threshold and kinetics. Consistent with our interpretation that, in male mice, there is a stronger gating mechanism to filter incoming signals during the active period, we discovered that the threshold for action potential firing was increased during the active period. These data suggest that layer 2/3 plPFC pyramidal neurons are not only more hyperpolarized during the light period, but are functionally more difficult to activate, requiring much larger depolarizations to elicit action potentials and relay information downstream. Though somewhat speculative, this could affect a wide range of behaviors, including emotionality, a notion supported by work demonstrating that pharmacological activation of plPFC neurons induces anxiogenic activity in mice, and acute stress enhances glutamatergic transmission in the PFC (10, 36, 38).

Neurophysiological sex differences in the PFC, and their respective behavioral outputs, have been documented and partly attributed to differences in synaptic signaling (22, 42). While exploring the effects of time-of-day on these fundamental properties of PFC cells, we fully embraced inclusion of both males and females, since use of both sexes (particularly inclusion of females) improves our overall understanding of brain function (43). While not designed explicitly as a sex-differences study, our results demonstrate that time-of-day did not strongly influence the function of PFC pyramidal neurons in female mice. This is consistent with the body of literature demonstrating sex differences in PFC function. For example, there are reported sex differences in synaptic function glutamate receptor expression and, in rats, higher basal release of glutamate in other layers of the PFC (44, 45). There are also sex differences in response to environmental and pharmacological stressors (46). When compared to male rats, female rats can display a lower threshold for impaired working memory after PFC injections of benzodiazepine inverse agonists that activate the stress system (47). Further, there are clear sex differences in mPFC dendritic growth, microglia activity, and astrocyte morphology in response to stress (48). We speculate that if the underlying mechanisms that mediate information throughput and plasticity are fundamentally different in males and females, then time-of-day changes in information filtering may not be as crucial to optimal plPFC function in female mice.

Our findings support a cell endogenous mechanism underlying daily changes in the physiology of layer 2/3 plPFC pyramidal neurons. This prompted us to explore how time-of-day impacts intrinsic postsynaptic properties such as ionic currents and overall conductance. There was a large effect on the current density and conductance of pyramidal neurons in male mice. Specifically, when these neurons were hyperpolarized below the equilibrium potential for K^+^, we discovered that current density increased throughout the light period, peaking between the late light period and early dark period. This effect translated into an overall increase in conductance. Given that conductance was highest around the beginning of the active period, when these neurons are most hyperpolarized, we posited that this was due to an increased number of open K^+^ channels and the outflow of K^+^ cations. Consistent with this prediction, when we blocked K^+^ channel mediated currents (by replacing K^+^ with Cs^+^ in our internal recording solution), the time-of-day effect on current density and conductance was completely abolished at voltages near or below the K^+^ equilibrium potential. Although internal Cs^+^ was not sufficient to block the time-of-day effect on current density at depolarized voltage greater than the K^+^ equilibrium potential, it greatly reduced overall current density and conductance.

Circadian desynchronization has myriad consequences for whole-animal physiology. This ranges from peripheral effects on metabolic and immune responses, to cognitive deficits and altered sleep (13, 49, 50). Our own prior work showed ECD reduced apical dendrite complexity and impaired cognitive flexibility in male mice. Our present results demonstrate that the consequences of circadian desynchronization extend to the cell function level, reducing information throughput of plPFC neurons by increasing the threshold for action potential firing, which is critical for communication with other brain regions, such as the hippocampus and amygdala. GIRK channels play a demonstrated role in pyramidal neurons and interneurons throughout the infralimbic and prelimbic cortices (21–23, 51). Dynamic changes in GIRK channels underlie stress-induced deficits in cognitive flexibility, and decreased GIRK channel function can impair fear learning, likely through altered communication between the PFC and hippocampus (23, 52). Our data show that ECD increases GIRK channel activity, and we speculate this tips the balance of inter-region communication. Together, this is in agreement with previous studies demonstrating the relationship between GIRK channel function and cognitive flexibility, and demonstrates that increased GIRK channel sensitivity may serve as a substrate for the deficits in cognitive flexibility observed in ECD mice (23, 49, 52).

It is now well accepted that the mPFC is heterogeneous at the anatomical and physiological levels (12), and perhaps the temporal dimension needs to be included in this framework. We believe our work reveals that fundamental aspects of cell and circuit function can be impacted by time-of-day, and underlines the importance of understanding how daily rhythms influence neural function throughout the brain. Our findings underscore that if we are to build comprehensive models of cell-circuit-behavioral outputs, we must address the relevant experimental and biological variables that impact these circuits, including biological time.

## Methods

### Mouse model(s)

All animal procedures and experiments were approved by the University of Massachusetts Amherst Institutional Care and Use Committee in accordance with the U.S. Public Health Service Policy on Humane Care and Use of Laboratory Animals and the National Institutes of Health *Guide for the Care and Use of Laboratory Animals*. Male and female wild-type mice (Charles River, Wilmington, MA, USA) on a C57BL/6N background (10-16 weeks old) were used for these studies. All mice were group-housed in light-tight housing boxes at 25°C, under a 12:12-hr light:dark (LD) cycle, with food and water available *ad libitum*. For circadian desynchronization experiments, mice were transferred to a 10:10h LD cycle and acclimated for at least three weeks prior to experimentation. LD cycles in housing boxes were offset so that experiments from each zeitgeist time (ZT) bin occurred at the same external (real world) time each day. For electrophysiology studies mice were anesthetized in a chamber with isoflurane before euthanasia by decapitation.

### Brain slice electrophysiology

Two mice were simultaneously euthanized 1-hr prior to their ZT bin (i.e., mice were euthanized at ZT23 for recording bin ZT0-4). After euthanasia, brains were immediately removed and the forebrain was blocked while bathing in a 0-4°C oxygenated N-methyl-D-glucamine (NMDG) - 4-(2-hydroxyethyl)-1-piperazineethanesulfonic acid (HEPES) cutting solution composed of (mM): 92 NMDG, 2.5 KCl, 1.25 NaH_2_PO_4_, 30 NaHCO_3_, 3 sodium pyruvate, 2 thiourea, 20 HEPES, 10 MgSO_4_, 0.5 CaCl_2_, 25 glucose, 20 sucrose. Cutting solution was brought to pH 7.4 with ∼17mL of 5M HCl (53). The forebrains were mounted adjacent to each other and sectioned simultaneously on a vibratome (VT1200S, Leica Biosciences, Buffalo Grove, IL, USA) with a sapphire knife (Delaware Diamond Knives, Wilmington, DE, USA) yielding roughly three slices containing the PFC from each (250-μm) per mouse. Slices were transferred and allowed to recover for 30-45 min in room temperature recording artificial cerebrospinal fluid (aCSF) solution composed of (mM): 124 NaCl, 3.7 KCl, 2.6 NaH_2_PO_4_, 26 NaHCO_3_, 2 CaCl_2_, 2 MgSO_4_, 10 glucose. aCSF had a final pH of 7.3-7.4, osmolarity of 307-310 mOsmos, and was continuously bubbled using 95% 0_2_/5% C0_2_. For recordings, brain slices were transferred to a perfusion chamber containing aCSF maintained at 34-37°C with a flow rate of 1mL/min. Neurons were visualized using an upright microscope (Zeiss Axoskop 2, Oberkochen, Germany). Recording electrodes were back-filled with experiment-specific internal solutions as follows (mM): Current-clamp and voltage-clamp; 125 K-gluconate, 10 KCl, 10 NaCl, 5 HEPES, 10 EGTA, 1 MgCl_2_, 3 NaATP and 0.25 NaGTP (liquid-junction potential (LJP) = ∼14.5 mV; Predicted E_K_ = ∼-95 mV). Voltage-clamp cesium-based internal solution; 140 CsCl, 5 MgCl_2_, 1 EGTA, 10 HEPES, 3 NaATP, and 0.25 NaGTP (LJP = ∼4.2 mV). All internal solutions were brought to pH 7.3 using KOH or CsOH at 301-304 mOsm. Patch electrodes with a resistance of 3-5MΩ were guided to neurons with an MPC-200-ROE controller and MP285 mechanical manipulator (Sutter Instruments, Novato, CA, USA). Patch-clamp recordings were collected through a UPC-10 USB dual digital amplifier and Patchmaster NEXT recording software (HEKA Elektronik GmbH, Reutlingen, Germany). Current clamp voltage-step protocols were performed from the cell endogenous resting membrane potential and used 500ms 10-pA steps from −100 pA to +190 pA. Voltage clamp current-step protocols were performed from V_H_= −70 mV, and used 10-mV steps from −120 mV to +30 mV. All compounds were obtained from Tocris Cookson, Cayman Chemical, and Sigma Aldrich.

Individual recording locations were plotted (with neurons outside of the target area excluded from analysis) to qualitatively confirm an equal distribution of recording sites between ZT bins 0-4, 6-10, 12-16, and 18-22 (***SI Appendix*, Fig. S1 A-D**). The electrophysiological heterogeneity of PFC pyramidal neurons is well described. A small percentage (∼20%) of all recorded neurons had unique characteristics in resting membrane properties and action potential dynamics that were independent of time-of-day (hereafter Type II neurons; ***SI Appendix*, Fig. S2 A-H**). Here we define Type I neurons as non-burst firing pyramidal neurons that are quiescent at rest, have a resting membrane potential (RMP) < −65 mV, membrane resistance < 200 MΩ, antipeak amplitude > −10 mV, and a qualitatively low velocity (measured as ΔmV/ms). Most notably, compared to Type I (most abundant) neurons, Type II neurons (less abundant) displayed a much higher action potential velocity and hyperpolarizing antipeak (***SI Appendix*, Fig. S2 A**,**B**). They also had a more depolarized RMP and decreased action potential firing threshold (***SI Appendix*, Fig. S2 E**,**F**). These categorical Type II neurons are likely a combination of off-target non-pyramidal neurons, and a small population of phenotypical distinct pyramidal neurons. Due to these clear qualitative and quantitative differences independent of ZT bin, and that they represented a small proportion of recorded neurons, we excluded the far less abundant Type II neurons from analysis in our following experiments.

### Experimental design and statistical analysis

Only neurons with input resistance > 70 MΩ were studied. Neurons were not considered for further analysis if series resistance exceeded 50MΩ or drifted >10% during baseline. Rheobase was calculated as the first current step to elicit an action potential and action potential dynamics (threshold, decay tau, and half-width) were obtained from the first evoked action potential to avoid variance in ion channel function due to repeated action potential firing. G*Power 3.0 software (Franz Faul, Uni Kiel, Germany) was used to conduct our power analysis, for a *p* value of <0.05 with 90% power. Adequate sample sizes were based upon expected effect sizes from similar experiments. Raw data files were analyzed in the Patchmaster NEXT software or converted using ABF Utility (Synaptosoft) for analysis in MiniAnalysis (Synaptosoft). N-values for analysis and presented in figures represent individual cells. Where applicable, extreme outliers were identified using the ROUT method with a conservative Q = 0.5%. To control for biological variability between groups N = 4-8 mice per group was used. To control for within animal variability 2-3 brain slices were collected per mouse. For experiments including the use of drug, only one cell per slice was used. Statistical comparison of effects between each time-period was made using a full model two-way ANOVA (column, row, and interaction effects) unless otherwise noted. Statistics were calculated using Prism 9 (Graphpad Software, San Diego, CA, USA).

## Acknowledgements

We would like to acknowledge and thank Dr. James Peters at Washington State University for his invaluable input and discussions during the preparation of this manuscript. Illustrations were created using Biorender.com. This work was supported by a CAREER grant 1553067 by the National Science Foundation, and R01 DK119811 from the National Institutes of Health to I.N.K.

## References

1. E. R. Woodruff, et al., Coordination between Prefrontal Cortex Clock Gene Expression and Corticosterone Contributes to Enhanced Conditioned Fear Extinction Recall. eNeuro 5 (2018).

2. M. J. McCarthy, D. K. Welsh, Cellular Circadian Clocks in Mood Disorders: Journal of Biological Rhythms (2012) https://doi.org/10.1177/0748730412456367 (September 14, 2020).

3. M. Popoli, Z. Yan, B. S. McEwen, G. Sanacora, The stressed synapse: the impact of stress and glucocorticoids on glutamate transmission. Nature Reviews Neuroscience 13, 22–37 (2012).

4. F. Sotres-Bayon, C. K. Cain, J. E. LeDoux, Brain Mechanisms of Fear Extinction: Historical Perspectives on the Contribution of Prefrontal Cortex. Biological Psychiatry 60, 329–336 (2006).

5. E. K. Miller, J. D. Cohen, An Integrative Theory of Prefrontal Cortex Function. Annu. Rev. Neurosci. 24, 167–202 (2001).

6. Y. Kawaguchi, Y. Kubota, GABAergic cell subtypes and their synaptic connections in rat frontal cortex. Cereb Cortex 7, 476–486 (1997).

7. G. Radnikow, D. Feldmeyer, Layer- and Cell Type-Specific Modulation of Excitatory Neuronal Activity in the Neocortex. Front. Neuroanat. 12 (2018).

8. R. P. Vertes, Interactions among the medial prefrontal cortex, hippocampus and midline thalamus in emotional and cognitive processing in the rat. Neuroscience 142, 1–20 (2006).

9. R. Saffari, et al., NPY+-, but not PV+-GABAergic neurons mediated long-range inhibition from infra-to prelimbic cortex. Transl Psychiatry 6, e736 (2016).

10. E. Y. Yuen, et al., Acute stress enhances glutamatergic transmission in prefrontal cortex and facilitates working memory. PNAS 106, 14075–14079 (2009).

11. A. V. Zaitsev, N. V. Povysheva, G. Gonzalez-Burgos, D. A. Lewis, Electrophysiological classes of layer 2/3 pyramidal cells in monkey prefrontal cortex. J Neurophysiol 108, 595–609 (2012).

12. D. E. Moorman, M. H. James, E. M. McGlinchey, G. Aston-Jones, Differential roles of medial prefrontal subregions in the regulation of drug seeking. Brain Research 1628, 130–146 (2015).

13. I. N. Karatsoreos, S. Bhagat, E. B. Bloss, J. H. Morrison, B. S. McEwen, Disruption of circadian clocks has ramifications for metabolism, brain, and behavior. PNAS 108, 1657–1662 (2011).

14. B. E. Kalmbach, D. H. Brager, Fragile X mental retardation protein modulates somatic D-type K+ channels and action potential threshold in the mouse prefrontal cortex. J Neurophysiol 124, 1766–1773 (2020).

15. W.-K. Deng, X. Wang, H.-C. Zhou, F. Luo, L-type Ca2+ channels and charybdotoxin-sensitive Ca2+-activated K+ channels are required for reduction of GABAergic activity induced by β2-adrenoceptor in the prefrontal cortex. Mol Cell Neurosci 101, 103410 (2019).

16. E. R. Workman, et al., Rapid antidepressants stimulate the decoupling of GABA(B) receptors from GIRK/Kir3 channels through increased protein stability of 14-3-3η. Mol Psychiatry 20, 298–310 (2015).

17. B. Bano-Otalora, et al., Daily electrical activity in the master circadian clock of a diurnal mammal. Elife 10, e68179 (2021).

18. L. M. Hablitz, H. E. Molzof, J. R. Paul, R. L. Johnson, K. L. Gamble, Suprachiasmatic nucleus function and circadian entrainment are modulated by G protein-coupled inwardly rectifying (GIRK) channels. J Physiol 592, 5079–5092 (2014).

19. K. I. van Aerde, D. Feldmeyer, Morphological and Physiological Characterization of Pyramidal Neuron Subtypes in Rat Medial Prefrontal Cortex. Cerebral Cortex 25, 788–805 (2015).

20. C. Piette, et al., Intracellular Properties of Deep-Layer Pyramidal Neurons in Frontal Eye Field of Macaque Monkeys. Front Synaptic Neurosci 13, 725880 (2021).

21. E. M. Anderson, S. Demis, H. D’Acquisto, A. Engelhardt, M. Hearing, The Role of Parvalbumin Interneuron GIRK Signaling in the Regulation of Affect and Cognition in Male and Female Mice. Front Behav Neurosci 15, 621751 (2021).

22. E. M. F. de Velasco, et al., Sex differences in GABABR-GIRK signaling in layer 5/6 pyramidal neurons of the mouse prelimbic cortex. Neuropharmacology 95, 353–360 (2015).

23. E. M. Anderson, et al., Suppression of pyramidal neuron G protein-gated inwardly rectifying K+ channel signaling impairs prelimbic cortical function and underlies stress-induced deficits in cognitive flexibility in male, but not female, mice. Neuropsychopharmacology 46, 2158–2169 (2021).

24. K. Kaufmann, et al., ML297 (VU0456810), the First Potent and Selective Activator of the GIRK Potassium Channel, Displays Antiepileptic Properties in Mice. ACS Chem. Neurosci. 4, 1278–1286 (2013).

25. D. Chaudhury, L. M. Wang, C. S. Colwell, Circadian regulation of hippocampal long-term potentiation. J Biol Rhythms 20, 225–36 (2005).

26. D. H. Loh, et al., Misaligned feeding impairs memories. eLife 4, e09460 (2015).

27. J. R. Paul, et al., Circadian regulation of membrane physiology in neural oscillators throughout the brain. European Journal of Neuroscience 51, 109–138 (2020).

28. L. Chrobok, et al., Daily changes in neuronal activities of the dorsal motor nucleus of the vagus under standard and high-fat diet. J Physiol (2021) https://doi.org/10.1113/JP281596.

29. L. McMartin, M. Kiraly, H. C. Heller, D. V. Madison, N. F. Ruby, Disruption of circadian timing increases synaptic inhibition and reduces cholinergic responsiveness in the dentate gyrus. Hippocampus 31, 422–434 (2021).

30. P. Xu, A. Chen, Y. Li, X. Xing, H. Lu, Medial prefrontal cortex in neurological diseases. Physiological Genomics 51, 432–442 (2019).

31. T. Otsuka, et al., Adverse Effects of Circadian Disorganization on Mood and Molecular Rhythms in the Prefrontal Cortex of Mice. Neuroscience 432, 44–54 (2020).

32. D. A. Bechtold, J. E. Gibbs, A. S. Loudon, Circadian dysfunction in disease. Trends Pharmacol Sci 31, 191–8 (2010).

33. Y. Hou, et al., Long-term variable photoperiod exposure impairs the mPFC and induces anxiety and depression-like behavior in male wistar rats. Exp Neurol 347, 113908 (2022).

34. J. H. Harkness, et al., Diurnal changes in perineuronal nets and parvalbumin neurons in the rat medial prefrontal cortex. Brain Struct Funct 226, 1135–1153 (2021).

35. N. K. Wallace, F. Pollard, M. Savenkova, I. N. Karatsoreos, Effect of Aging on Daily Rhythms of Lactate Metabolism in the Medial Prefrontal Cortex of Male Mice. Neuroscience 448, 300–310 (2020).

36. B. Li, L.-L. Chang, K. Xi, Neurotensin 1 receptor in the prelimbic cortex regulates anxiety-like behavior in rats. Progress in Neuro-Psychopharmacology and Biological Psychiatry 104, 110011 (2021).

37. K. A. Petrie, et al., The Neurotensin Agonist PD149163 Increases Fos Expression in the Prefrontal Cortex of the Rat. Neuropsychopharmacol 29, 1878–1888 (2004).

38. A. Saitoh, et al., Activation of the prelimbic medial prefrontal cortex induces anxiety-like behaviors via N-Methyl-D-aspartate receptor-mediated glutamatergic neurotransmission in mice. Journal of Neuroscience Research 92, 1044–1053 (2014).

39. M. L. Phillips, H. A. Robinson, L. Pozzo-Miller, Ventral hippocampal projections to the medial prefrontal cortex regulate social memory. Elife 8, e44182 (2019).

40. A. R. Fusilier, et al., Dysregulated clock gene expression and abnormal diurnal regulation of hippocampal inhibitory transmission and spatial memory in amyloid precursor protein transgenic mice. Neurobiol Dis 158, 105454 (2021).

41. J. Yan, et al., Short-term sleep deprivation increases intrinsic excitability of prefrontal cortical neurons. Brain Research 1401, 52–58 (2011).

42. S. Andrade, et al., Effect of progesterone on the expression of GABA(A) receptor subunits in the prefrontal cortex of rats: implications of sex differences and brain hemisphere. Cell Biochem Funct 30, 696–700 (2012).

43. R. M. Shansky, A. Z. Murphy, Considering sex as a biological variable will require a global shift in science culture. Nat Neurosci 24, 457–464 (2021).

44. C. J. Perry, E. J. Campbell, K. D. Drummond, J. S. Lum, J. H. Kim, Sex differences in the neurochemistry of frontal cortex: Impact of early life stress. Journal of Neurochemistry 157, 963–981 (2021).

45. J. I. Pena-Bravo, R. Penrod, C. M. Reichel, A. Lavin, Methamphetamine Self-Administration Elicits Sex-Related Changes in Postsynaptic Glutamate Transmission in the Prefrontal Cortex. eNeuro 6, ENEURO.0401-18.2018 (2019).

46. E. Y. Yuen, J. Wei, Z. Yan, Estrogen in prefrontal cortex blocks stress-induced cognitive impairments in female rats. J Steroid Biochem Mol Biol 160, 221–226 (2016).

47. R. M. Shansky, et al., Estrogen mediates sex differences in stress-induced prefrontal cortex dysfunction. Mol Psychiatry 9, 531–538 (2004).

48. J. L. Bollinger, I. Salinas, E. Fender, D. R. Sengelaub, C. L. Wellman, Gonadal hormones differentially regulate sex-specific stress effects on glia in the medial prefrontal cortex. J Neuroendocrinol 31, e12762 (2019).

49. G. L. Pearson, M. Savenkova, J. J. Barnwell, I. N. Karatsoreos, Circadian desynchronization alters metabolic and immune responses following lipopolysaccharide inoculation in male mice. Brain Behav Immun 88, 220–229 (2020).

50. D. J. Phillips, M. I. Savenkova, I. N. Karatsoreos, Environmental disruption of the circadian clock leads to altered sleep and immune responses in mouse. Brain Behav Immun 47, 14–23 (2015).

51. E. M. Anderson, S. Demis, B. Wrucke, A. Engelhardt, M. C. Hearing, Infralimbic cortex pyramidal neuron GIRK signaling contributes to regulation of cognitive flexibility but not affect-related behavior in male mice. Physiol Behav 242, 113597 (2021).

52. N. C. Victoria, et al., G Protein-Gated K+ Channel Ablation in Forebrain Pyramidal Neurons Selectively Impairs Fear Learning. Biol Psychiatry 80, 796–806 (2016).

53. J. T. Ting, et al., Preparation of Acute Brain Slices Using an Optimized N-Methyl-D-glucamine Protective Recovery Method. JoVE (Journal of Visualized Experiments), e53825 (2018).

