## Supplemental Figures and legends for "Circadian desynchronization attenuates information throughput of prefrontal cortex pyramidal neurons in mice"

**This PDF file includes:**

Figures S1 to S2

1. **Supporting Figures**

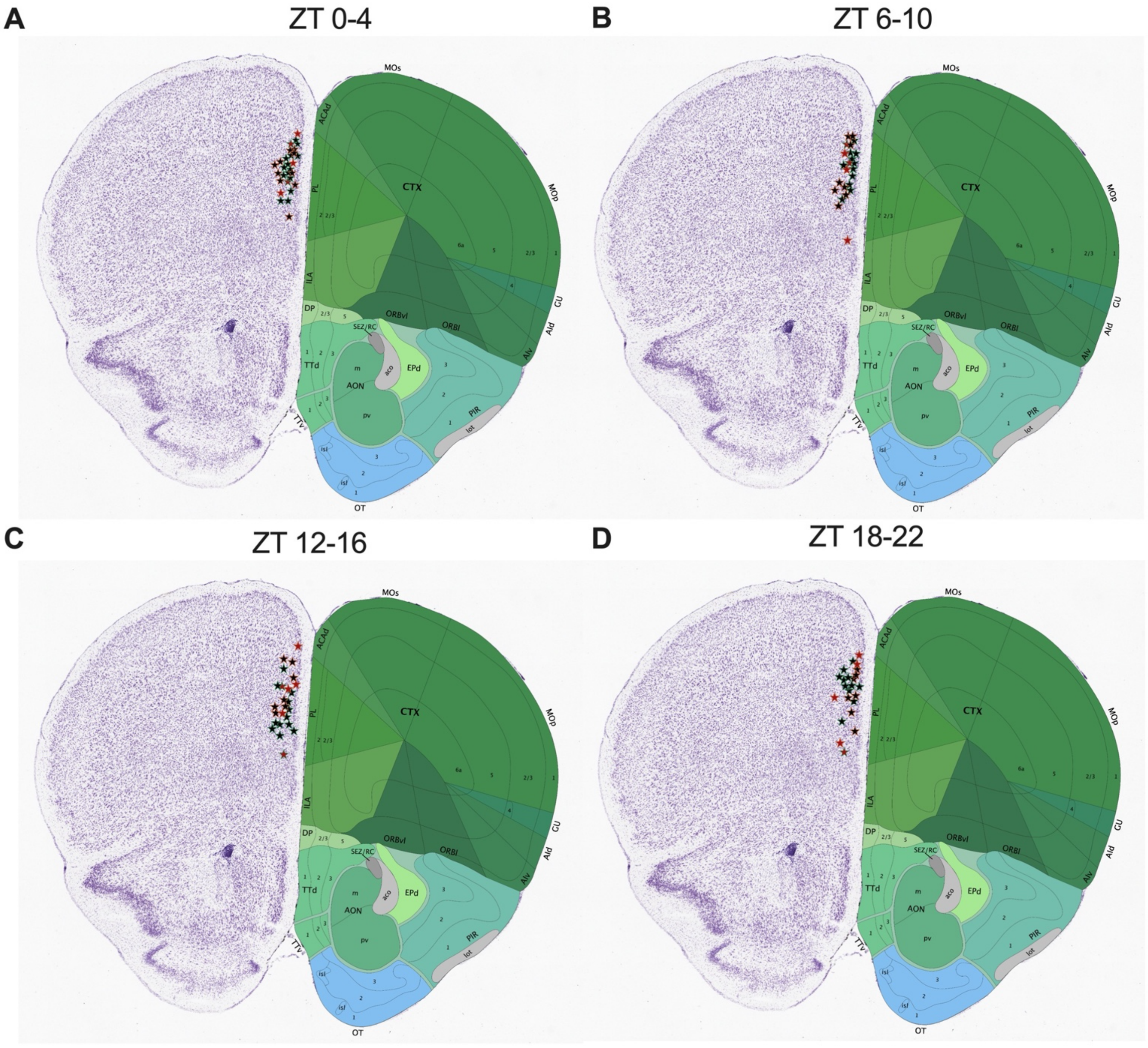

**Fig. S1.** *Recording map for layer 2/3 plPFC pyramidal neurons.* Coronal sections of forebrain showing individual recording sites from majority of neurons that were imaged at **(A)** ZT0-4, **(B)** 6-10, **(C)** 12-16 and **(D)** 18-22 for basal membrane property, sEPSC, and evoked action potential experiments in male (*bluish green outline*) and female (*vermillian outline*) mice. Stars filled with black represent 'Type I' neurons included for analysis and red stars represent Type II/III neurons excluded from analysis.

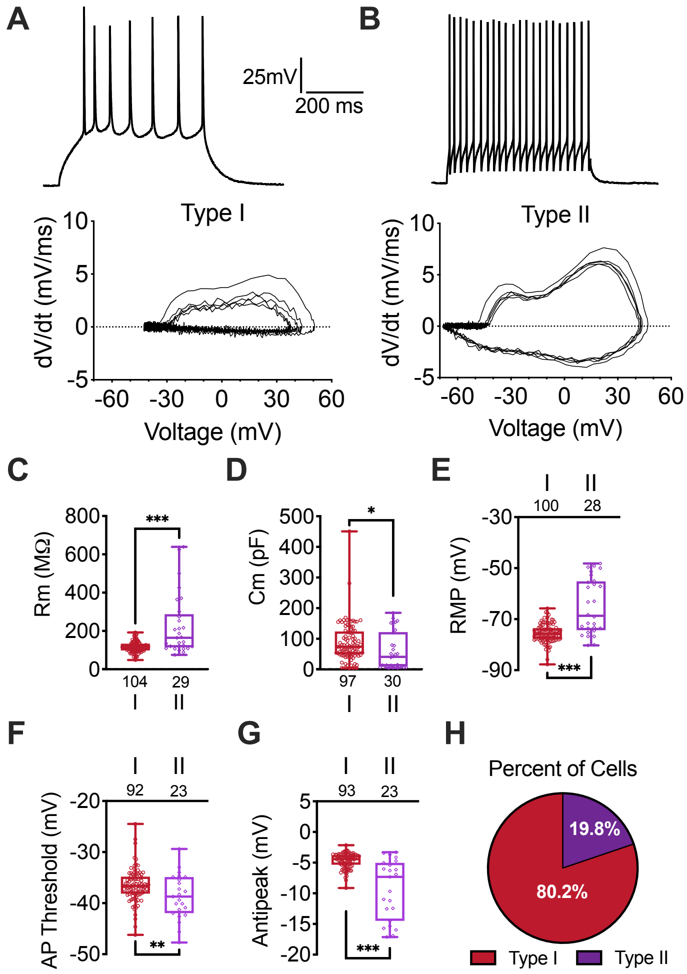

A
B
25mV
200 ms
Type I
Type II
dV/dt (mV/ms)
Voltage (mV)
C
D
E
Rm (MΩ)
Cm (pF)
RMP (mV)
F
G
H
AP Threshold (mV)
Antipeak (mV)
Percent of Cells
19.8%
80.2%
Type I
Type II

**Fig. S2.** *Categories and distinct physiological characteristics of plPFC neurons.* **(A)** Representative evoked action potential traces and phase plot diagram of first five action potentials (*bottom*) illustrating differences in velocity, trajectory, and amplitude in Type I and **(B)** Type II neurons. **(C)** Boxplot comparison of membrane resistance (Rm; t = 6.656, df = 131; $p$ <0.001), **(D)** membrane capacitance (Cm; t = 2.003, df = 125; $p$ = 0.05), **(E)** resting membrane potential (RMP t = 8.133, df = 126; $p$ <0.001), **(F)** action potential (AP) threshold (t = 2.674, df = 113; $p$ = 0.009), **(G)** antipeak amplitude (t = 8.27, df = 114; $p$ < 0.001), and **(H)** Percentage of recorded cells displaying Type I or Type II characteristics (combined among all ZT bins and sexes; calculated by *n* values from AP threshold). Unpaired student t-test, * $p< 0.05$, ** $p< 0.01$, ***$p< 0.001$.
